# Optimal reaching subject to computational and physical constraints reveals structure of the sensorimotor control system

**DOI:** 10.1101/2023.05.14.540714

**Authors:** Patrick Greene, Amy Bastian, Marc H. Schieber, Sridevi V. Sarma

## Abstract

Optimal feedback control provides an abstract framework describing the architecture of the sensorimotor system without prescribing implementation details such as what coordinate system to use, how feedback is incorporated, or how to accommodate changing task complexity. We investigate how such details are determined by computational and physical constraints by creating a model of the upper limb sensorimotor system in which all connection weights between neurons, feedback, and muscles are unknown. By optimizing these parameters with respect to an objective function, we find that the model exhibits a preference for an intrinsic (joint angle) coordinate representation of inputs and feedback and learns to calculate a weighted feedforward and feedback error. We further show that complex reaches around obstacles can be achieved by augmenting our model with a path-planner based on via points. The path-planner revealed “avoidance” neurons that encode directions to reach around obstacles, and “placement” neurons that make fine-tuned adjustments to via point placement. Our results demonstrate the surprising capability of computationally constrained systems and highlight new characteristics of the sensorimotor system.

## Introduction

Optimal feedback control (OFC) has emerged in the last two decades as a useful and general theoretical framework for understanding the sensorimotor system [TJ02; Sco04; Sco+15]. In OFC, the motor cortex is viewed through the lens of control theory: the motor cortex and effectors such as the arm are viewed as dynamical systems, where the motor cortex is a controller whose job is to move the effector in a way that optimally achieves some goal, such as reaching a target in minimal time or energy use. The controller also receives sensory feedback, both visual and proprioceptive, which is used to estimate the state of the effector and compare it to the desired state in order to continually generate corrections to the current movement path.

### OFC explains features of actual movements, but is agnostic on implementation

OFC has been successful in explaining many features of movement such as the shape of reach trajectories and velocity profiles [FMB13], triphasic patterns of neural activity and changes in directional tuning [TMK07], and biases in the preferred directions of neurons [LS13]. However, while providing a useful framework for the overall structure of the sensorimotor system, it is agnostic on the details of how feedback control may be implemented by populations of neurons, although recent studies have sought to tease out the brain regions that may form the components of an OFC system [Tak+21; DAl+22]. Under dynamical systems based approaches like OFC, the activity of a region or population of neurons may not consistently represent something like the velocity of a hand or the coordinates of a target [Vya+20; Chu+10].

### Many experiments have found neural correlates of coordinate systems

This representational neutrality leaves us the job of understanding how the neural correlates of coordinate systems and their transformations fit into the more abstract OFC framework. Experiments using wrist movement have shown that subsets of M1 neurons display activity that modulates with the preferred direction of muscle as well as the direction of movement [KHS99]. In a reaching task, Kurata found that neurons could be classified into types whose activity depended on target locations in head-centered visual coordinates or motor coordinates, as well as neurons that had either differential or non-differential activity for both coordinates and thus could be contributing to coordinate transformations [Kur07]. Most directly, [WH06; WH07] compared the mutual information between the firing rate of primary motor, dorsal pre-motor, and ventral pre-motor (PMv) neurons and the movement direction during a reach task in three different coordinate systems: an extrinsic cartesian, intrinsic joint angle, and intermediate shoulder-centered system that rotates with the shoulder. They found evidence for all three coordinate systems, with a small bias for a shoulder-centered system in M1 and for both shoulder-centered and joint angle coordinate systems in PMv.

### Optimizing the sensorimotor architecture for simple reaches

In the first part of this paper, we ask what aspects of the motor control system can be uncovered within the OFC framework when we simulate realistic effector kinematics and apply computational limitations in the controller during simple reaches. In particular, we seek to gain insight into the following questions: 1) Is there a preferred coordinate system in the representation of movement? and 2) How are feedback and input signals incorporated by an optimal controller into its dynamics?

We study these questions by creating an architecture wherein the motor cortex is modeled as a linear system that provides outputs to the end effector, a nonlinear physics-based model of the primate arm, and the state of the effector is fed back to the motor cortex. The connection weights between neurons, muscles, and feedback are all unknown. We give the controller input and feedback in both cartesian and joint angle coordinate systems, and then optimize its neural connectivity over a large set of reaches with respect to a simple objective function that penalizes position error and energy use. *Importantly, the controller is able to make accurate and efficient reaches between any two points in the workspace and is not fit to experimental data or optimized for a particular task such as a center-out reach*. We find that a strong preference emerges for joint angle coordinates and that the system naturally learns to compute an error feedback signal.

### Augmenting the optimal sensorimotor architecture for complex reaches

In the second part of this paper we consider how more complex reaches around obstacles can be achieved by augmenting the control system that was developed for simple reaches. We explore the use of via points -intermediate targets that guide the reach along an arbitrary path -as inputs to our M1 model and show that, when carefully chosen, they allow the model to successfully navigate around obstacles. We propose a simple method for determining via points that maintains a high rate of obstacle avoidance and show that it can be implemented by a (non-linear) feed forward neural network that serves as a PM-like “planning area” for our simple reach model with linear controller. We use lesion studies to analyze groups of neurons in the trained network and find that some neurons tend to specialize for calculating via points in particular regions of space or reach directions, while others are broadly responsible for biasing the via point to the left or right of the line between the initial position and target.

## Online Methods

### Arm model

We constructed a two-dimensional, physics-based arm model with two joints and six muscles. There are two bones, an upper arm bone mimicking the humerus and a lower arm bone mimicking the ulna. Masses and lengths of the upper and lower arm segments were set to typical values of the Rhesus macaque (*Macaca mulatta*), taken from [CS00]. The joints consist of a shoulder and elbow joint, each of which has its angular motion soft-constrained by an exponentially increasing opposing force to realistic ranges (between -45 and 135 degrees for the shoulder, where 0 has the arm pointing out to the right, and between 0 and 135 degrees for the elbow, where 0 is complete extension).

Joint torques in the arm model are produced by a set of six muscles whose role is to mimic the action of one or more real muscle groups. The muscles consist of two uniarticular shoulder muscles, two uniarticular elbow muscles, and two bijoint muscles which span both the shoulder and elbow as shown in **Fig. 1A**. Each pair of muscles consists of a flexor and an extensor. Muscle moment arms were taken from [TMK07]. For simplicity, muscles are assumed to have linear dynamics of the form 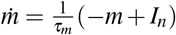 where *m* is the muscle activation, *τ*_*m*_ = 0.02 sec is the activation (and deactivation) time constant, and *I*_*n*_ is the neural input function which will be made explicit below. For a given muscle activation level, the torque about the spanned joint is calculated by multiplying the muscle activation by the maximum muscle force and the moment arm. The kinematics and equations of motion for the arm were derived using standard techniques (see i.e. [TMK07; LS13]). The equations of motion will be denoted 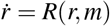, where 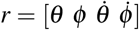 is the vector of joint angles and angular velocities.

**Figure 1:**
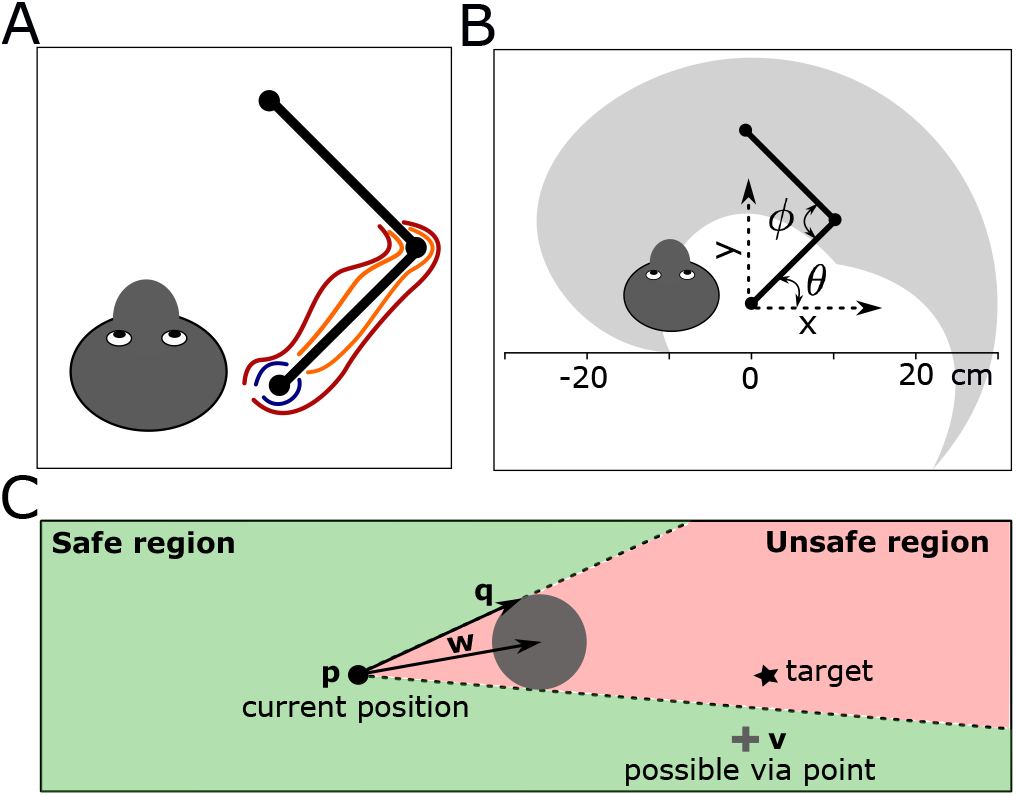
A) The model macaque arm, depicted with head to indicate forward direction and relative position and size. Three pairs of muscles are included in the model: shoulder (blue), elbow (orange), and bijoint (red) flexors and extensors. B) Joint angle coordinates *θ* and *ϕ* represent the angle between the upper arm and the *x* axis and the angle between the upper and lower arm, respectively. The crescent-shaped shaded gray region shows the reachable area of the arm. C) Illustration of the cone method for determining potential via point locations when an obstacle (shown as a gray circle) is present in the path of the reach. Via points may be placed anywhere in the safe region defined as the complement of the cone formed by the current position and the edge of the obstacle.

### Neural model

The arm model is controlled using a linear dynamical system with feedback shown on the first line of equation 1. The set of neuronal units has some firing rate at time *t* denoted by the vector *x*(*t*), and all units receive input from other neuronal units with weights given by the neuron-neuron connectivity matrix *A* and a bias vector *B*_0_ that sets the baseline firing rates. The *i, j* entry of *A* is the weight that unit *i* places on input from unit *j*. In addition, units receive a set of inputs denoted by Σ_*i*_*B*_*i*_*u*_*i*_, where each *u*_*i*_ is a static input vector to the system and *B*_*i*_ is the corresponding weight matrix that maps the input into the neural state space. We consider two different coordinate systems in the horizontal plane -an orthogonal cartesian coordinate system that is fixed with respect to the body and represents points as (*x, y*) pairs with the origin at the shoulder, and a joint angle coordinate system that represents points as (*θ, ϕ*) pairs consisting of the shoulder angle relative to the *x* axis and the elbow joint angle as shown in **Fig. 1B**. The inputs *u*_1_ and *u*_2_ are the starting position of the hand in cartesian and joint angle coordinates, respectively, and inputs *u*_3_ and *u*_4_ are the position of the target in cartesian and joint angle coordinates. Finally, the neuronal units receive feedback denoted by Σ_*i*_*C*_*i*_*v*_*i*_(*t*), where each *v*_*i*_(*t*) is a time-varying feedback function and *C*_*i*_ is the corresponding weight matrix. *v*_1_(*t*) and *v*_2_(*t*) are the current hand position in cartesian and joint angle coordinates, and *v*_3_(*t*) is the muscle activation level for each of the 6 muscles.

The neural model is coupled to the muscle dynamics via the neural input function *I*_*n*_ = *σ* (*D*_0_ + *D*_1_*x*) where *D*_1_ is the neural population to muscle connectivity matrix, *D*_0_ is a bias term, and *σ* (*x*)= 1/(1 + *e*^*− x*^) maps the input between 0 and 1. In summary, the dynamics of our coupled motor cortex and arm model are given by:

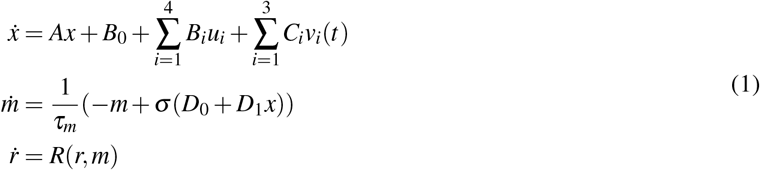

and illustrated by the diagram in **Fig. 2**. Smoothly time-varying noise was added to the joint angle feedback in the form of a draw from a gaussian process with a standard deviation (amplitude of the gaussian kernel function) ranging from 1.8 degrees to 36 degrees and autocorrelation scale (standard deviation of the kernel function) of 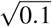. The autocorrelation scale sets how rapidly the noise changes over time. In our case, we have it change relatively slowly so that it represents a drift in the estimated joint angles.

**Figure 2:**
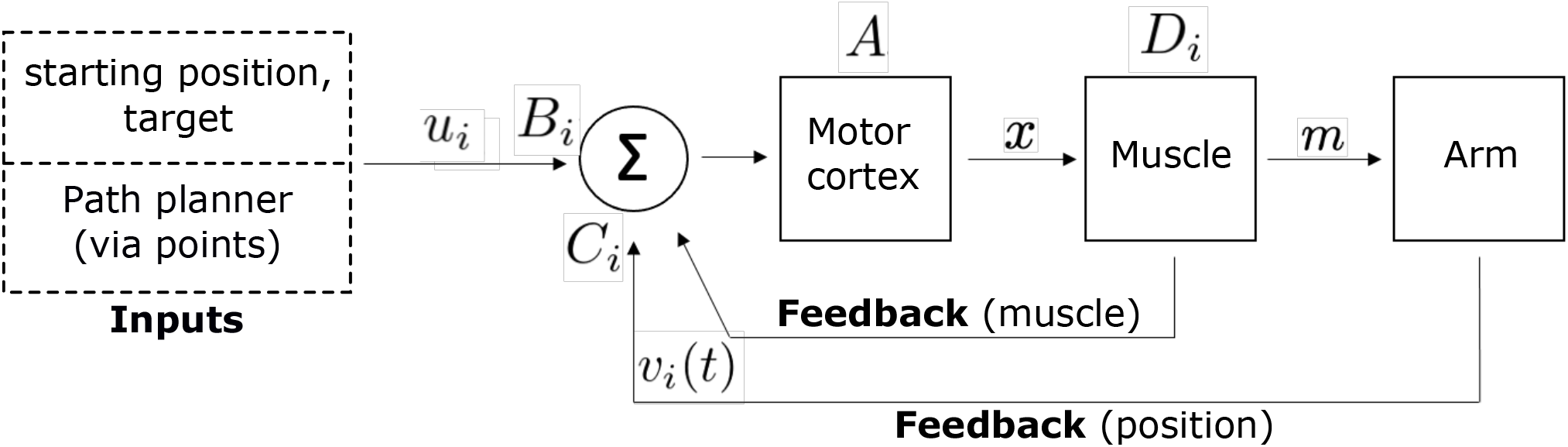
A block diagram of the motor cortex and arm model.

Because each neural unit in our model has a smooth output that represents a mean-subtracted firing rate and can have both positive and negative connections to other units, we consider the units as abstractly representing neurons or populations of neurons with similar sets of inputs and activation patterns. We therefore use the terms ”neurons” or ”neural units” to refer to them.

### Optimization

Given a training set of *S* samples consisting of starting points and targets (no obstacles) spread uniformly over the reachable space and a time window of length *T* in which to make each of the reaches, we optimized the matrices *A*, {*B*_*i*_}, {*C*_*i*_}, *D*_0_, *D*_1_ simultaneously in our model with respect to the following objective function:

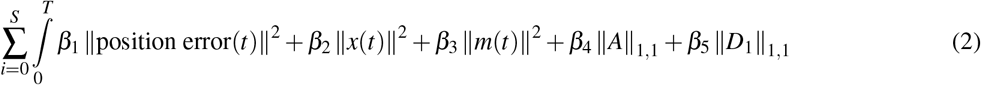

where position error(*t*) is the time-varying difference between the position of the hand and the target, *β*_1_, *β*_2_, and *β*_3_ are fixed parameters which set the amount of penalization for position error, neural activity, and muscle activity, and *β*_4_ and *β*_5_ set the penalization on neuron to neuron and neuron to muscle connectivity. We set *β*_2_ and *β*_3_ approximately 1/200 the size of *β*_1_, and *β*_4_ and *β*_5_ approximately 1/10^5^ the size of *β*_1_. Note that the norms for the *A* and *D* matrices are the entry-wise *L*^1^ norms (the sum of absolute values of the entries of the matrix) to promote sparsity, while the norms for the other terms are the standard *L*^2^ norm squared (sum of squares of entries). *T* was set to 1 second and *S* = 10, 000 training samples were used. We set the number of neurons in our model (the dimension of *x*) to 20. Experiments with varying numbers of neurons showed that performance was relatively consistent across a wide range between 10 and 200 neurons thanks to the sparsity penalty on *A*, which effectively pruned unneeded neurons. The optimization was accomplished using Pontryagin’s adjoint sensitivity method [Pon+63], in which gradients are calculated by formulating and numerically solving an ordinary differential equation (ODE) called the adjoint ODE backwards in time. The system of differential equations comprising the equations of motion, neural and muscle dynamics, and adjoint equations was coded in python using the NumPy and SciPy numerical libraries and integrated using an adaptive fifth order Runge-Kutta method [Har+20; Vir+20]. Gradient descent using the BFGS algorithm with the calculated gradient vectors was performed until convergence [Vir+20].

### Obstacles

Reaches around obstacles were accomplished by augmenting the linear neural controller described above with time-varying intermediate targets (“via points”) that replace the fixed final target input used in unobstructed reaches. Based on the assumption that the optimized linear controller makes reaches in approximately straight lines, we propose a simple method, called the visual cone method, for calculating via points that allow the effector to move around an obstacle. At any given time, the set of rays passing from the current hand position to the set of points that produce trajectories that intersect the obstacle forms a cone, and any via point outside this cone is in what we call the “safe region,” meaning that a reach to a point in this region will never hit the obstacle if it travels in a straight line from the starting point to the via point. The safe region and its cone-shaped complement, the ”unsafe region”, for a given starting point and target are shown in **Fig. 1C**. Finding the cone and whether a via point is safe is a simple calculation. Assume we know the current hand position **p**, the location of the center of the obstacle a distance *d* from **p**, and the obstacle’s radius *r*. Let **w** denote the vector from the current position to the obstacle center and **v** the position of a potential via point. The vector **q** pointing along one side of the cone is found by rotating **w** by an angle *α* = *sin*^−1^(*r*/*d*). The potential via point **v** lies in the safe region if

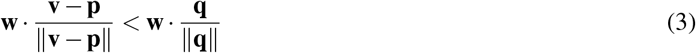

Physically, the safe and unsafe regions are closely related to whether a point is visible or not. The closest safe point is then found by projecting the target onto the nearest wall of the cone. Since trajectories of the model are in reality not exactly straight, we improve the obstacle avoidance rate by moving perpendicularly outward from the walls of the cone in both directions in small increments until we find a safe via point. This requires ”testing” points by running the forward model simulation; for the sake of speed we limit this to no more than 10 points on either side of the cone, although typically only one or two tests are needed as we accept the first valid point. The accepted via point is then fed to the linear controller as a new target, with updates occurring every 50 ms.

### Neural networks

Although the visual cone method was constructed with simplicity and biological feasibility of the computations in mind, we further investigate its feasibility by training feedforward neural networks on the via points generated by the cone method in order to determine how complex a network would need to be in order to reproduce the outputs of the method. To simplify the task, we trained the neural networks on fixed length reaches of 18 cm, with 6 cm diameter circular obstacles placed at the midpoint of each reach. The reach direction and region of the workspace in which the reach occurred were random, as was the shift of the obstacle relative to the centerline of the reach (the straight line between the starting point and target), which could be up to 2 radii in either direction. Rather than continuously predicting via points throughout the reach, the networks were only required to output the first via point, as this was the primary via point that allowed the reach to move around the obstacle. After 50 ms, all subsequent via points were simply set to the final target position. The neural networks were feedforward networks with ReLU activation, either 10, 20, 50, 100, or 200 neurons in each hidden or input layer (the 2 neuron output layer predicted the via point coordinates), and between 1 and 6 hidden layers. The networks were constructed using the PyTorch framework and trained for 1000 epochs with the Adam optimizer at a learning rate of 5×10^−4^ [Pas+17]. The most accurate network was then chosen for virtual lesion analysis. We clustered neurons in the final hidden layer of this network based on the vector of weights associated to each neuron (the set of weights that the neuron applies to inputs from the previous layer). This was accomplished by first calculating principal components and taking the top 3 as a reduced dimensional representation. K-means clustering was then used to identify clusters within the principal component space, with only consistently identified clusters across multiple random initializations and choices of cluster number *k* subject to analysis [Vir+20].

## Results

### Simple reaches: The optimized model reproduces several established features of the sensorimotor system

After optimizing our model, we find that our model is able to reproduce several important aspects of M1 neurons and their generated movement from the experimental literature as shown in **Fig. 3**. The neurons in our model develop tuning curves and direction preferences that do not change with the length of the reach, reproducing experimental results by [Geo+82; FSE93]. **Fig. 3C** shows that direction preferences do change when the reach occurs in a different part of the workspace, with the preferred direction rotating clockwise as the task region (shown by dashed boxes) moves from left to right, reproducing results by [CJU90]. Finally, the overall wrist velocity curve of the optimized model is bell-shaped, with longer reaches developing higher peak velocities in similar proportion to that seen in experiments [CD19].

**Figure 3:**
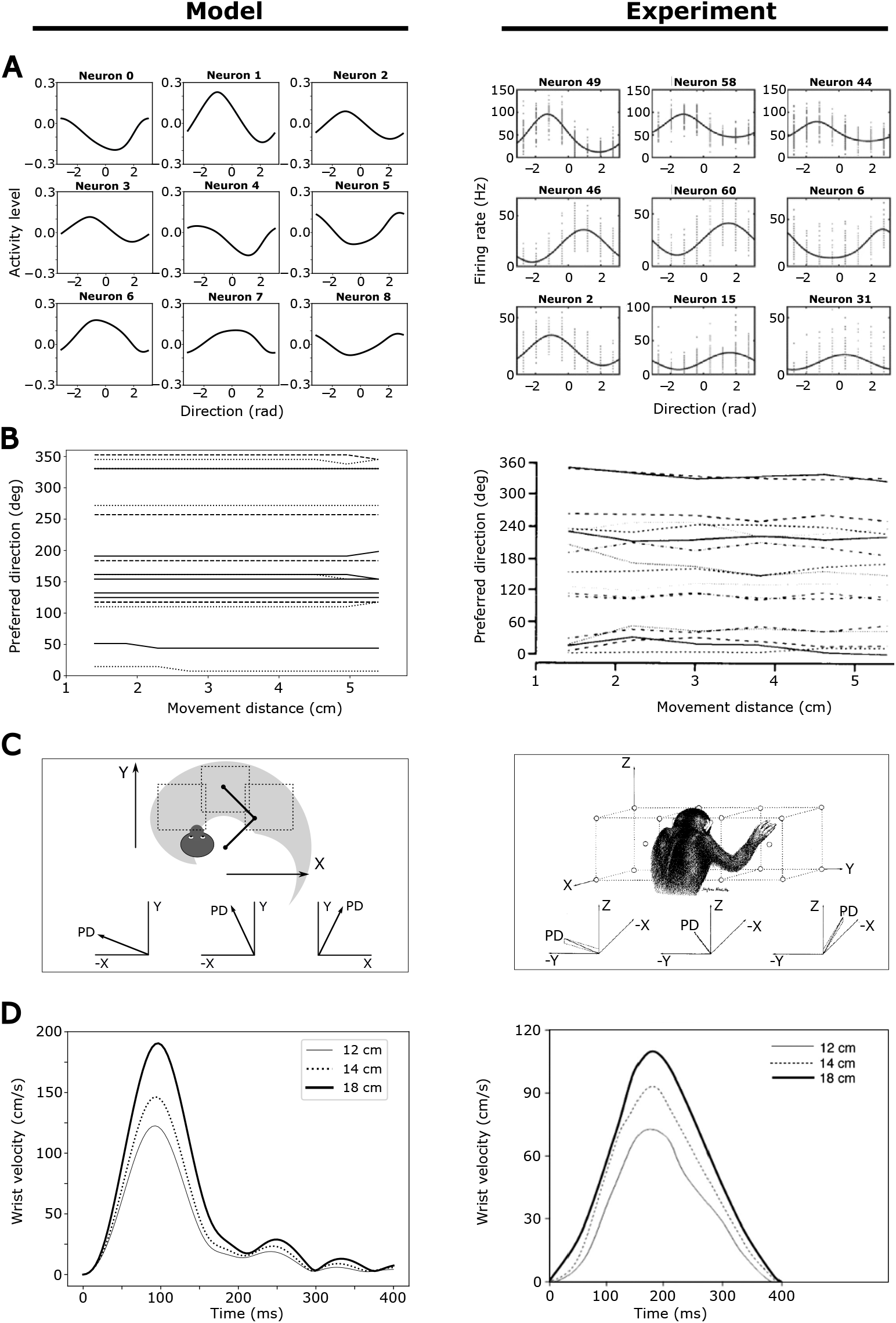
Comparison of model characteristics after optimization (left column) compared to experimental literature (right column). A) Tuning curves displayed by model neuron units on a center-out task, versus experimental tuning curves from [Tam+19] (plotted from data in [Mat+18]). B) Stability of neuron direction preferences across reach lengths in model versus experiments in [FSE93]. C) Rotation of the preferred directions of neurons as the workspace shifts from left to right. The region in which the center-out task was performed are indicated by the dashed squares (left, center, and right) in the model and by the boxes in the experimental illustration [CJU90]. The vector labeled ”PD” in both images indicates the preferred direction of a single neuron. D) Bell-shaped velocity curves for reaches to a target of varying length versus experimental data from [CD19].

### The controller weights joint angle input and feedback over cartesian coordinates

Optimization of the model occurs over a large set of random reaches throughout the workspace, and therefore it can perform a reach from any given starting point to any target. The center-out trajectories shown in **Fig. 4A** demonstrate that the model has learned an effective control policy that results in relatively straight and accurate reaches. In terms of the model parameters, the most apparent outcome of the optimization process is the difference in the sizes of the learned weights between the cartesian and joint angle input and feedback – the matrices *B*_1_ vs *B*_2_ (these weight the starting position in cartesian and joint angles, respectively), *B*_3_ vs *B*_4_ (weighting the target position), and *C*_1_ vs *C*_2_ (weighting the hand position), as shown in **Fig. 4B**. In all cases, the absolute values of the learned weight matrices corresponding to joint angles are more than 3 times larger on average than the learned weights for the corresponding cartesian coordinates. Accounting for differences in the scale of the coordinates increases this even further.

**Figure 4:**
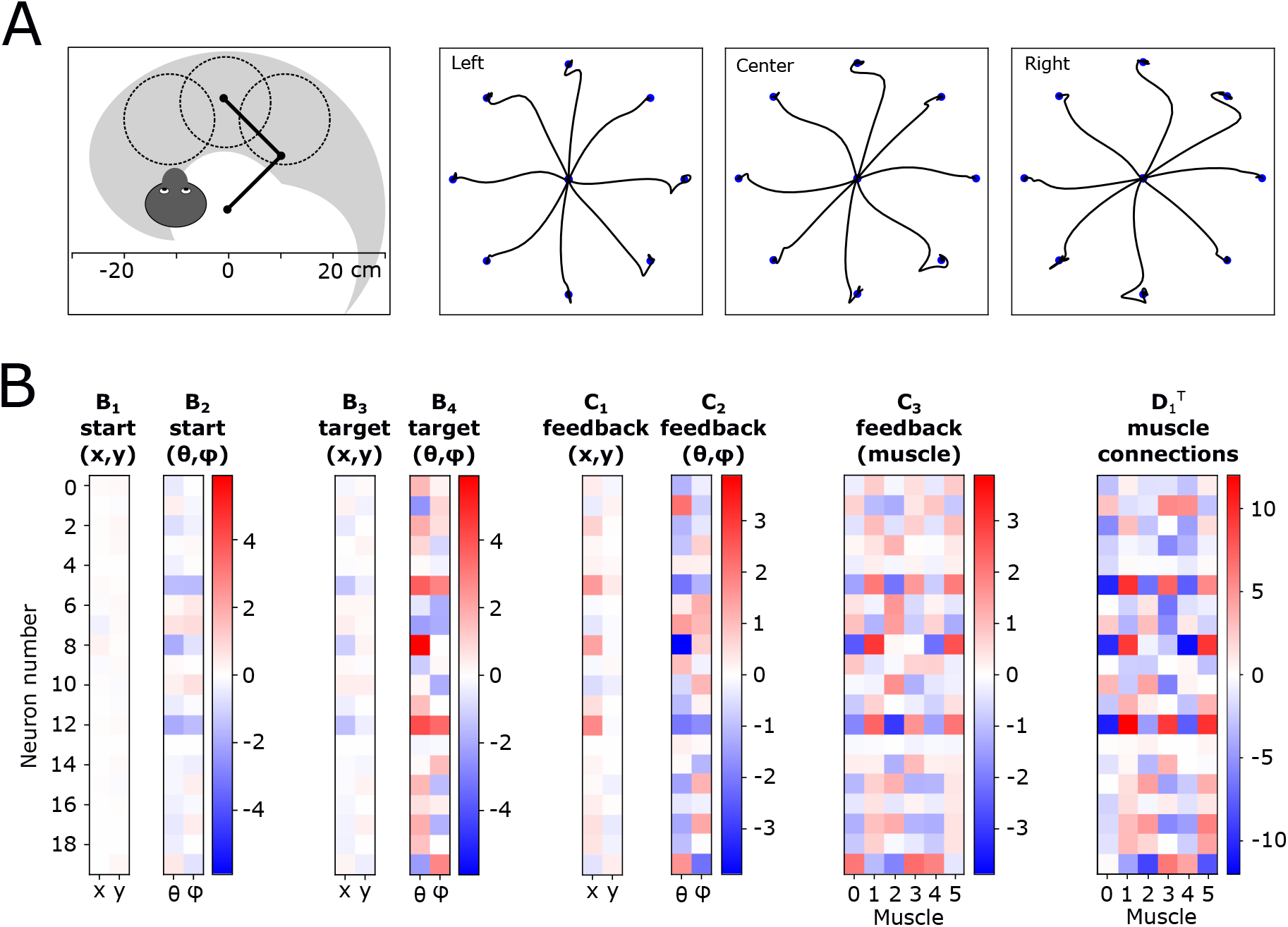
Cartesian versus joint angle weighting after optimization. A) Example center-out reaches of the optimized model in three different regions of the reachable space. In the leftmost figure, the reachable space is shown by the gray crescent-shaped area, and the dashed circles indicate the left, center, and right regions in which the center-out task was performed. B) Weight matrices after optimization.

### Feedback noise changes the operating mode of the controller rather than the coordinate system

Another important outcome of the optimization is that the feedback (*C*_2_) and start position (*B*_2_) matrices become scaled, negative versions of the target (*B*_4_) weight matrix, showing that the model naturally learns to compute both feedback (target minus feedback) and feedforward (target minus starting position) error signals. When we increase the noise level in the joint angle feedback and retrain the model, we find that, as expected, the weighting applied to the now unreliable joint angle feedback decreases, as shown in **Fig. 5A**. However, rather than compensating with increased levels of cartesian feedback, the weighting applied to the cartesian feedback remains relatively constant while the weight given to the joint angle starting position increases significantly. Further, the complexity of the controller, as shown by the density of the neuron-neuron connectivity matrix in **Fig. 5B** increases significantly. Rather than switch coordinate systems, these changes indicate that the model prefers to continue using joint angle coordinates, but now in a manner more strongly driven by the feedforward error.

**Figure 5:**
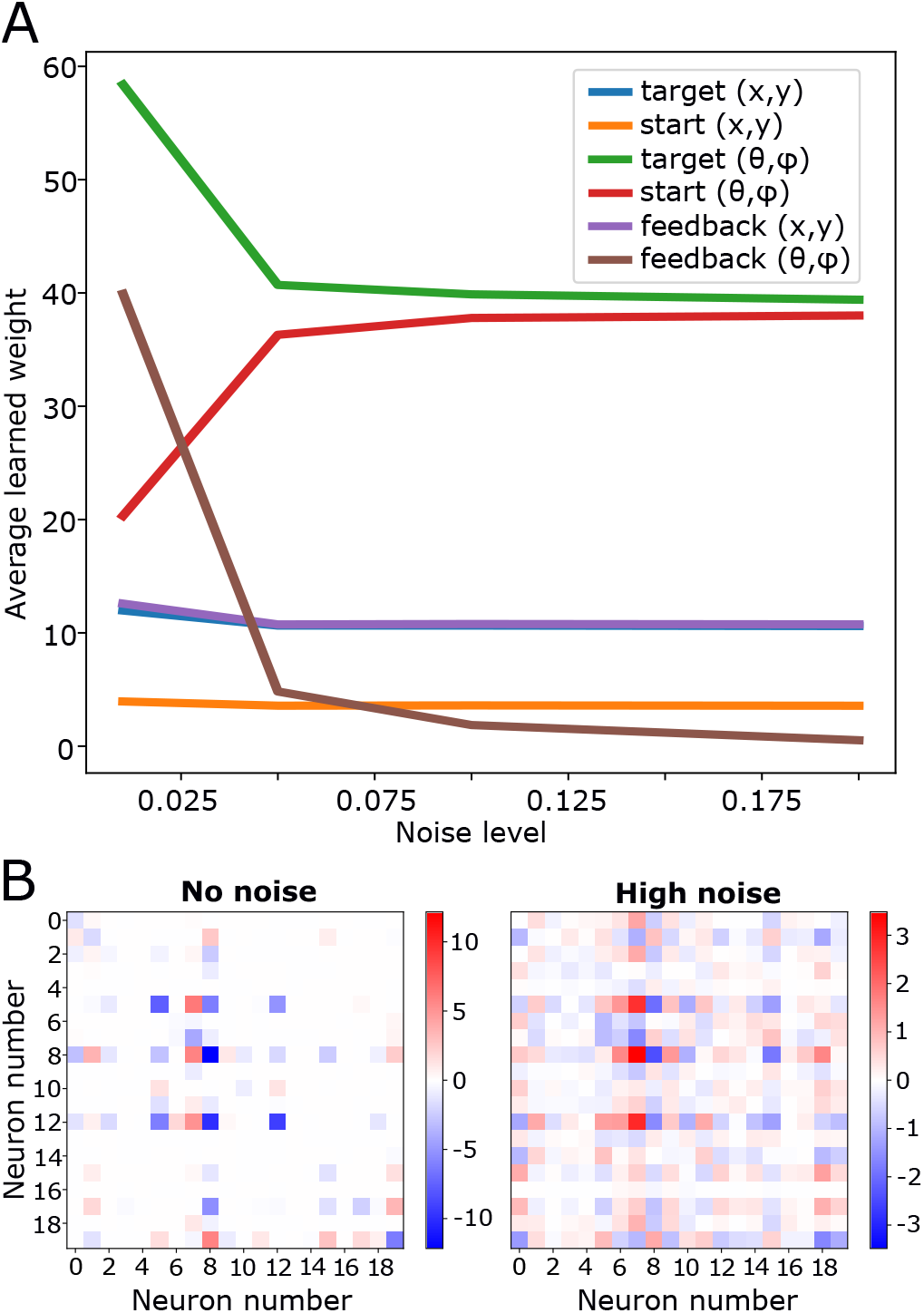
The effect of noise in joint angle feedback. A) Average absolute value of weights in each of the input and feedback matrices (shown in figure 4 as a function of the joint angle feedback noise level. B) Comparison of the neuron-neuron connectivity matrix in no noise versus high noise conditions.

### Complex reaches: The visual cone method provides a computationally simple means of finding via points

Although our linear neural system is effective at controlling the arm in simple reaches, we find that it struggles to navigate around an obstacle in a realistic way. We believe this is because there are two important nonlinearities which must be accounted for, as shown in **Fig. 6A**. The first is that when the center of the obstacle moves from one side of the centerline of the reach to the other, the direction of the reach must jump to the opposite side of the obstacle because this is now the shorter path. In addition, when the obstacle has shifted so much that it is no longer in the way, the shift in the reach must go to zero and remain there for any further shift of the obstacle. Our visual cone method takes into account these nonlinearities, and by using it to compute via points we are able to avoid hitting the obstacle 95% of the time.

**Figure 6:**
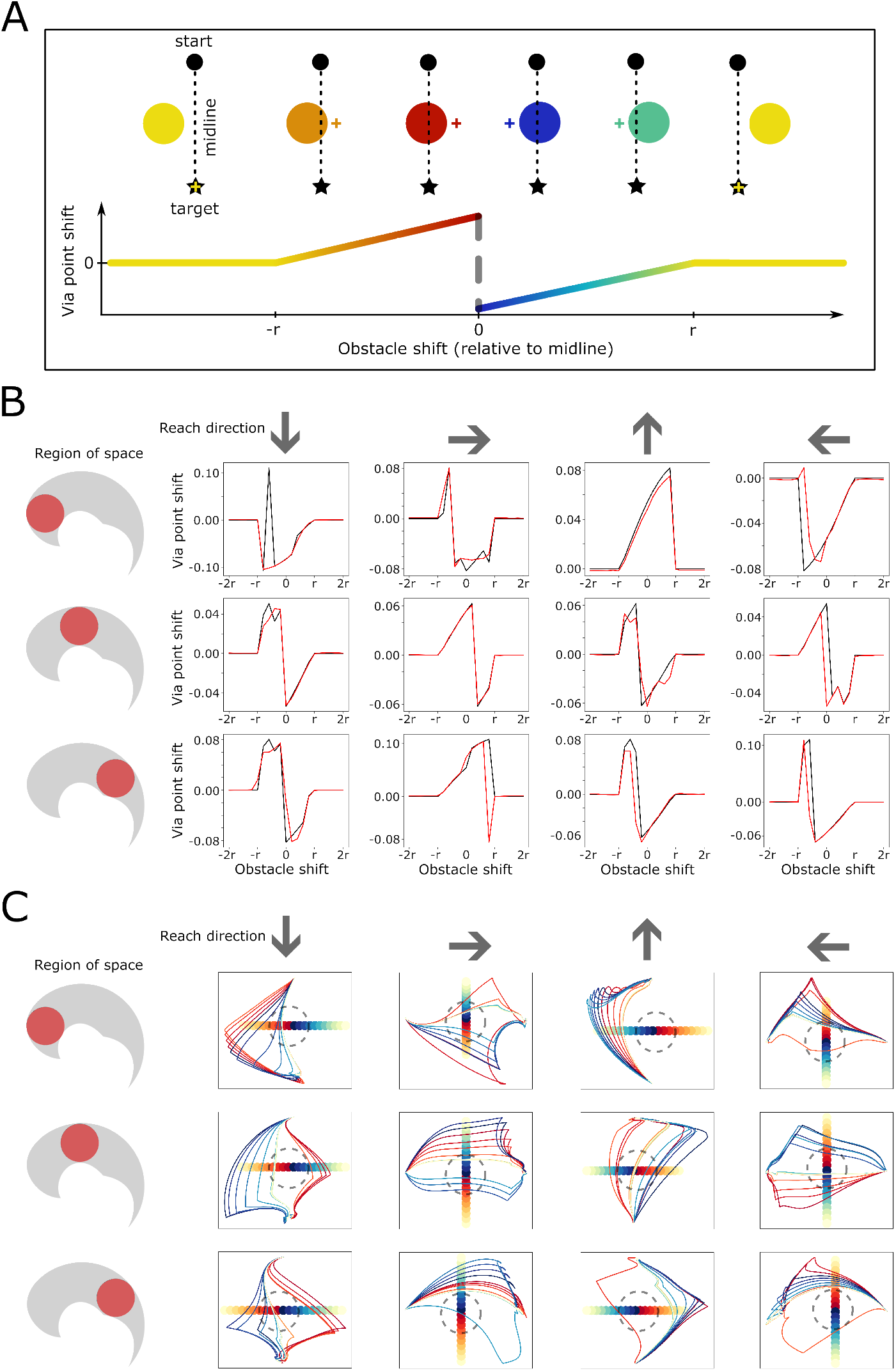
A neural network can learn cone-based via point placement. A) Illustration of how via point placement changes nonlinearly as an obstacle moves across the centerline of the reach. B) Comparison between the neural network’s via point placement (red) and the cone method (black). Each subplot shows the distance of via points from the centerline of a reach as a function of the obstacle position relative to the centerline. Each row of subplots shows a reach in a particular region of the workspace (highlighted in red in the graphic to the left) and each column is a particular reach direction (indicated by the arrows at the top of each column). C) Reach trajectories. Each subplot shows the trajectories of the set of reaches corresponding to the obstacle shifts indicated by the colored circles. Each colored circle indicates the position of the center of the obstacle, and the line of the same color shows the corresponding reach trajectory. The dashed gray circle indicates the actual size of the obstacle. As in B), rows and columns of the subplots indicate regions of space and reach directions, respectively.

### A neural network can learn the visual cone method

We fit a range of feedforward neural networks to via points generated by the visual cone method and found that a 3 hidden layer, 200 neuron network most accurately reproduced test set via points generated by the cone method, although most networks with at least 100 neurons and 2 hidden layers had satisfactory performance (see Supplemental Material Fig. S2A). **Figure 6B** compares the via point placement of the cone method (in black) versus the best trained neural network (in red) in 3 different regions of space and 4 reach directions. At the center of each region of space shown by the red circles on the leftmost column of **Fig. 6B**, an obstacle was placed and shifted by small increments up to 2 radii perpendicular to the centerline. For each obstacle shift, reaches were taken from one side of the circle to the other in the indicated directions. We see that the neural network is generally able to reproduce the visual cone method’s via points, and that the via point shift function often takes the expected zigzag shape shown in **Fig. 6A**. Individual reaches in each region of space and reach direction are shown in **Fig. 6C**. Each obstacle shift is colored according to the corresponding reach, and we can see how reaches typically move around the obstacle in the appropriate direction before reorienting towards the target. Due to the fact that the linear controller does not make perfectly straight reaches, i.e. for a given reach it may be biased to move toward the left or right, reaches sometimes take a longer path than necessary to accommodate this bias by going around the “wrong” side of the obstacle.

### Clustering reveals coarse ”avoidance” and fine-tuned ”placement” groups of neurons

When we cluster neurons in the final hidden layer of our path planning network based on their input connectivity (the weights that they apply to inputs from neurons in the previous layer), lesioning reveals two main classes of neurons. The first are those that seem to encode a rough direction in which to go around an obstacle and have an influence across all regions of space, which we refer to as obstacle ”avoidance” type neurons. Pre and post-lesion effects of this cluster are shown in **Fig. 7A** and **B**. The gray panels in A) indicate the spatial regions and reach directions where lesioning had a significant (*>* 50% relative difference) impact on the via point distance from the reach centerline. Lesioning this cluster produced a very pronounced shift of the via points across nearly all regions and reach directions while keeping the shape of the placement curves relatively unchanged. The direction of the post-lesion shift depended on the reach direction and spatial region, with a rightward shift for upward or downward reaches in the left and center regions (the red line appears shifted in different directions for the upward and downward reaches because the orientation is flipped; in the physical space the shifts are both to the right), and a leftward shift in the right region. A similar phenomenon occurs in the center versus left regions for leftward and rightward reaches. In contrast, a cluster of the second type consists of what we call ”placement” type neurons, which are those that have a limited spatial-directional influence, making more fine-tuned adjustments to the via point placement in specific regions of space and reach directions. As shown in **Fig. 7C** and **D**, these type of neurons play an important role in adjusting the via point away from the centerline for certain obstacle positions while leaving the rest of the via point placement curve unchanged. In this example, the cluster adjusts via point placement for obstacles that are in the 90 degree clockwise direction from the reach, but only in the center and right regions of space and rightward and upward reach directions.

**Figure 7:**
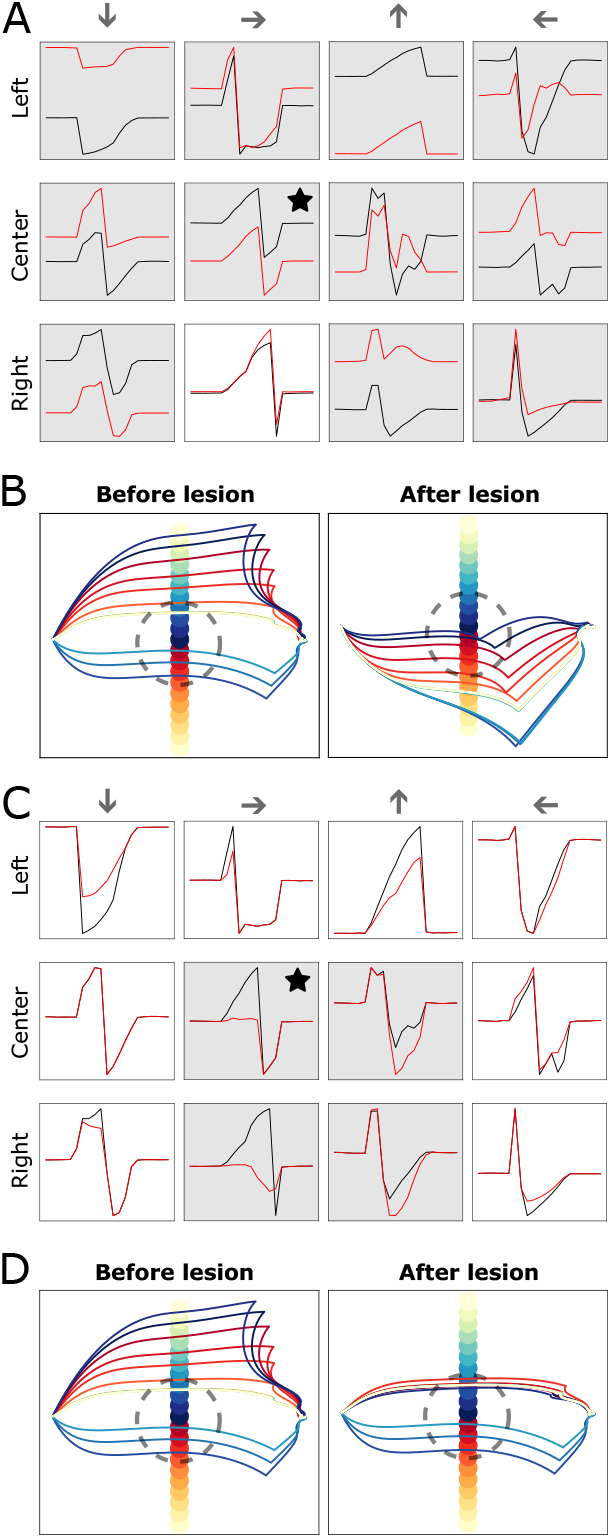
Effect of lesions in the neural network. A) Via point shift as a function of obstacle shift before (black) and after (red) a lesion of all neurons in the ”avoidance” cluster. Subplots with greater than 50% relative difference between pre and post-lesion are shaded in gray. B) Reach trajectories before and after the lesion corresponding to the starred subplot in A). C) Via point shift as a function of obstacle shift before (black) and after (red) a lesion of all neurons in a ”placement” cluster. D) Reach trajectories before and after the lesion corresponding to the starred subplot in C).

## Discussion

In optimizing the controller, we emphasize again that the model is not fit to an experimental data set and is not trained for a specific task such as center-out reaches. The final controller is not necessarily optimal for any particular reach, but rather is locally optimal within the class of linear controllers with respect to the average error over the entire training set of reaches. A single set of neural weights is required to give appropriate controller dynamics for all reaches. Despite this, the optimization process nearly always yields a solution that works well in controlling the arm across the entire reachable space regardless of initial weights and a wide range of tuning parameters.

Because they can be computed directly from muscle lengths, joint angles are more closely related to a coordinate representation that might be computed from proprioceptive feedback in the form of stretch and load information from muscle afferents. In contrast, extrinsic coordinates are more closely related to visual input and intermediate retinotopic coordinates. In primates, proprioceptive feedback is likely to be noisier than visual feedback, but this is balanced by the fact that visual feedback has much longer delays – approximately 135 ms versus 60 ms for the long-latency muscle response generated through the transcortical pathway involving M1 and 30 ms for spinal level feedback [Car81; Mat91]. When we increased the noise in the joint angle feedback but not the cartesian coordinate feedback during training, the model continued to use joint angle coordinates in a feedforward mode rather than switching to cartesian coordinates. This suggests that differences in noise level result in changes in the type of computation (feedforward versus feedback) rather than changes in the preferred coordinate system. Our model assumes no feedback delays in either coordinate system; adding significantly higher delays in the visually-inspired cartesian coordinate system would only further increase the preference toward joint angles.

If we only give the model access to cartesian coordinate input and feedback, we find that the linear controller is incapable of producing accurate and efficient reaches regardless of how much we penalize position error or muscular effort (see Supplemental Material Fig. S1B). We believe the model prefers to use joint angle coordinates because of the simpler error feedback relationship between muscle activations and the movement of the hand. A positive difference between the target and current joint angle always requires flexor activation while a negative difference always requires extensor activation, whereas in cartesian coordinates, the muscles required to correct a displacement in either the x or y coordinate may be completely reversed depending on where the hand is in space.

There are many possible coordinate systems, and in this study we chose two examples that are on opposite ends of the intrinsic/extrinsic spectrum. The important thing from the point of view of the controller is that the preferred coordinate system have a relatively direct relationship to the error, rather than that it be joint angles specifically. The fact that a simple linear controller is sufficient for a wide range of movements shows that controller complexity and cost due to the number of synapses and neurons that need to be maintained can be significantly reduced when the right coordinate representation is used. Although motor cortex is not necessarily a linear system, evidence indicates that this may be a reasonable approximation. Studies by Evarts, Cheney, and Fetz showed that many M1 neurons have activity that closely tracks the magnitude of applied torque across joints, and over a large part of the tested range, M1 neuron responses varied linearly with the level of static force [Eva68; CF80]. However, later work by Georgopoulos and many others using center-out tasks pointed to cartesian movement directions as the information encoded by M1 neurons [Geo+82]. Our model shows that cartesian direction preferences and cosine tuning curves are seen in center-out tasks even when computation occurs primarily in a joint angle coordinate system. Although in theory a sufficiently complex system could allow any coordinate system to be used, computational costs imposed by biology make it likely that coordinate representations are carefully chosen in the brain to account for the degrees of freedom and musculature of the arm, even if they are not as simple as the pure joint angle coordinates used in this study. In our model, we note that a coordinate system is not explicitly represented in the activity of the neurons themselves. After multiplication by the weight matrices, the neurons work with an abstract N-dimensional linear combination of the two joint angle coordinates, and their activity reflects muscle activation in our model. Despite the lack of an explicit representation in neural activity, the properties of the coordinate system in which inputs and feedback are presented to the controller are integral to its performance.

Our model learns to compute an error signal consisting of the difference between the current and target hand position. Several studies support the idea that an error signal is formed in M1 and that the size of the response increases with the error signal [Omr+16; IUK16]. This error feedback component of the control input allows for a simple controller because the error signal has a direct relationship to the output signal in the appropriate coordinate system. In contrast, in order to accomplish primarily feedforward control, as we see occurs in our model when feedback is noisy, the connectivity and therefore the complexity of the motor system must be greatly increased, as seen in **Fig. 5B**. This occurs because, without accurate error feedback, the controller must invert the dynamics of the effector in order to create a sequence of signals that produce a trajectory which reaches the target. Instead of simply reacting to an error signal, it must now “plan” in the sense of using the connections between neurons to produce dynamics in the neural state space that, when composed with the arm system, are the identity – the output of the system (the hand position) is equal to the input (the target). However, since the arm dynamics are non-linear, the computationally constrained linear brain model cannot accomplish this effectively and trajectories are degraded.

The neuron to muscle connections generally form an overcomplete basis for movement since there are 6 muscles but only 4 are required to move the arm to any position (an extensor and flexor for each degree of freedom). A priori, one might expect that an *L*^1^ penalty on neuron to muscle connections such as the one we impose on our model would result in a minimal basis with each neuron population activating only a single muscle. What we find instead is that each neuron population tends to connect to multiple muscles across both joints and simultaneously deactivate antagonist muscles, in a fashion similar to corticomotoneuronal muscle fields in primates.

For reaches around obstacles, the inherent nonlinearity of the task suggests that a more complex computation must occur with support from non-M1 regions, an idea backed up by lesion experiments [MK77]. Via points are a possible means by which a controller for simple reaches could be augmented to produce more complex reaches. Several studies have shown, using a double-step paradigm in which target locations are changed mid-reach, that online course-correction mechanisms of the type that would presumably be used in via point navigation result in smooth trajectories and depend on the posterior parietal cortex to integrate visual and somatosensory information and present this to PM and M1 [Arc+15; SM15]. Whether there exists a ”path planning system” that provides intermediate targets for a lower level system that accomplishes simple reaches in the way we have modeled is unknown. However, we believe that such a system may be biologically feasible, and our lesion study of a neural network implementation has provided potential neural tuning characteristics of such a system.

## Acknowledgements

S.V.S., M.H.S., and P.G. disclose support for the publication of this study from the National Institutes of Health (NIH), grant R01-NS110423.

## Author contributions

P.G. and S.V.S. conceived the hypothesis. P.G. developed the model and analysed the results with feedback from M.H.S., S.V.S., and A.B. P.G., S.V.S., and M.H.S. wrote the manuscript.

## Competing interests

The authors have no competing interests.

## Supplemental Material

**Supplemental Material, Figure S1:**
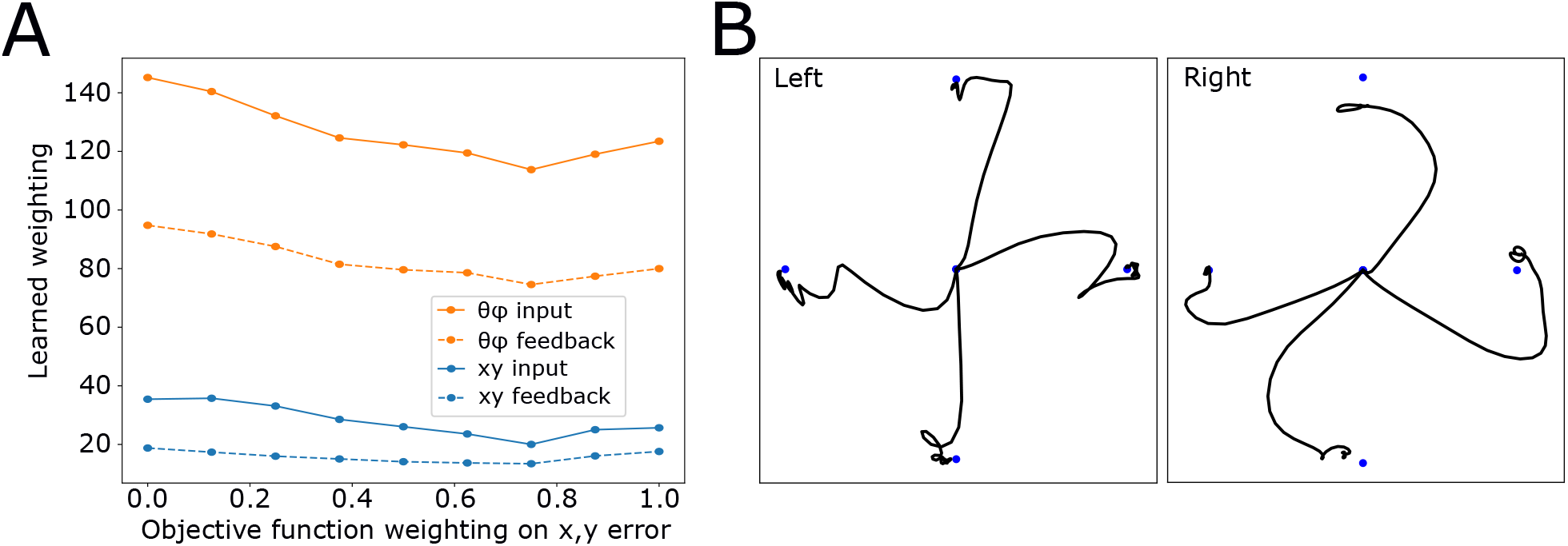
A) Total weighting on cartesian versus joint angle coordinates in input and feedback as a function of the relative weight of the cartesian position error. B) Example reaches in a model with only cartesian input and feedback.

To test whether the difference in coordinate weighting was due to the way that the objective function measured the position error, we modified it to include two position error terms, one measuring the sum of squares between the current hand position and the target in cartesian coordinates, and a second measuring the sum of squares of the angular error in joint angle coordinates. As shown in **Fig. S1A**, we found that varying the relative weighting of the two error terms did not change the preference for joint angle coordinates, with joint angle input and feedback weighting remaining more than 3 times larger even when only cartesian error was considered. Since a straight line is much more simply expressed in cartesian coordinates, one might expect that forcing the model to use only cartesian input and feedback would result in straighter reaches. However, when we remove all joint angle input and feedback and retrain the model from scratch, we find that it produces significantly less efficient, curvier reaches that sometimes do not even reach the target. Examples of typical center-out reaches from the fully cartesian model are shown in **Fig. S1B**.

**Supplemental Material, Figure S2:**
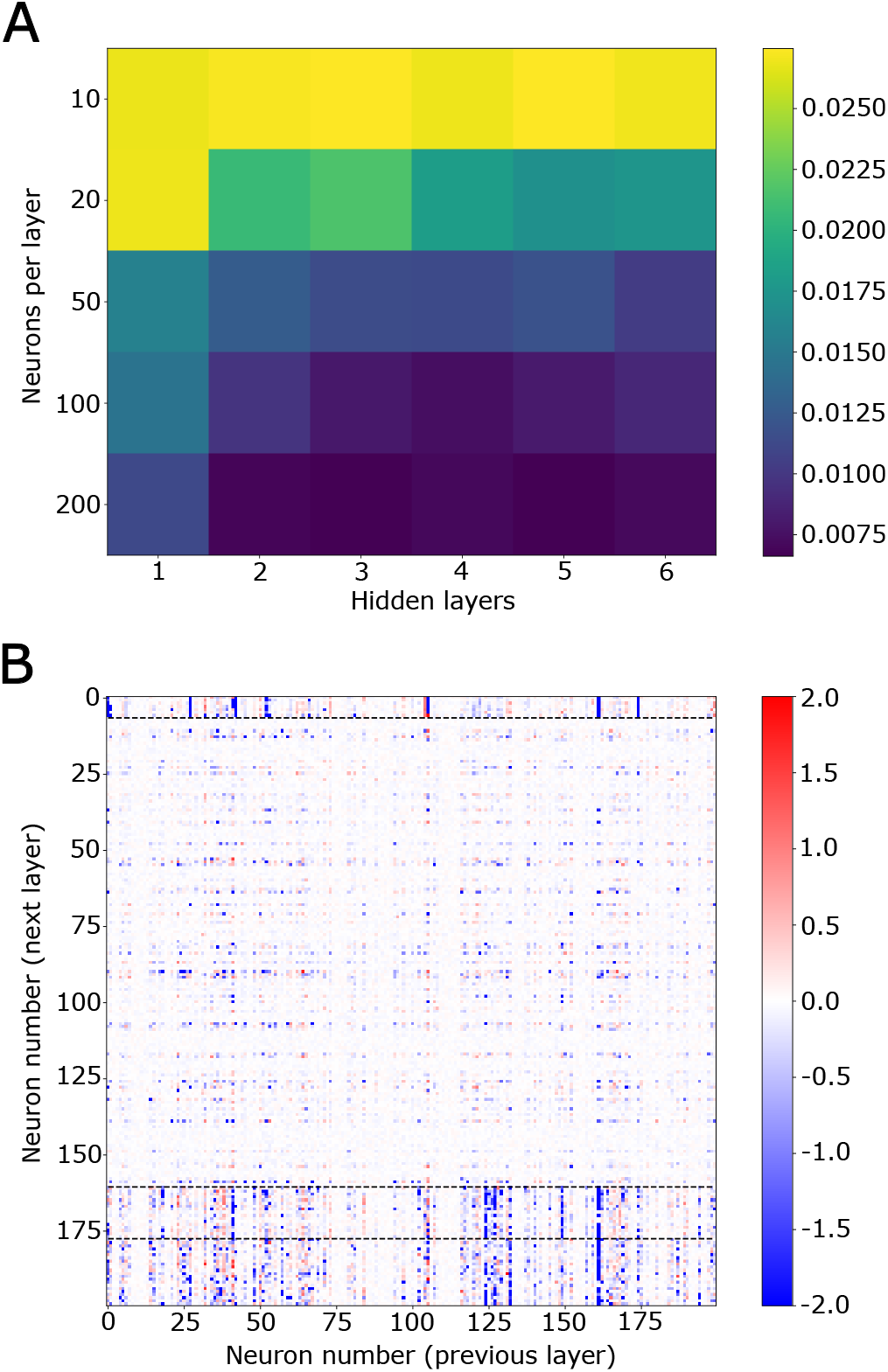
Via point neural network accuracy and clustering. A) Via point prediction accuracy on the test set for 30 feedforward neural networks with size ranging between 1 -6 hidden layers and 10 -200 neurons per hidden layer. B) Weight matrix for the final hidden layer of the neural network. Each row represents the weights that the final hidden layer applies to inputs from neurons in the previous layer. Clustering was performed on the rows of the weight matrix and the rows were reordered according to their cluster membership (indicated by dashed horizontal lines). From the top, clusters 1, 3, and 4 are examples of ”placement” clusters, while cluster 2 is an ”avoidance” cluster.

## Notes

### Competing Interest Statement

The authors have declared no competing interest.

